# PI3 kinase-unrelated effects of LY294002 and LY303511 on serotonin-induced Ca^2+^ and cAMP signaling

**DOI:** 10.1101/2024.05.05.592569

**Authors:** Polina D. Kotova, Ekaterina A. Dymova, Olga A. Rogachevskaja, Stanislav S. Kolesnikov

## Abstract

The phosphoinositide 3-kinase (PI3K) is involved in regulation of multiple intracellular processes. Although the inhibitory analysis is generally employed for validating a physiological role of PI3K, increasing body of evidence suggests that PI3K inhibitors can exhibit PI3K-unrelated activity as well. Here we studied effects of PI3K inhibitor LY294002 and its inactive analogue LY303511 on Ca^2+^ and cAMP signals initiated by serotonin. In the present study several monoclonal HEK293 cell lines were used, in particular, monitoring of Ca^2+^ signals were carried out on Fura-2 loaded cells expressed recombinant serotonin 5-HT_2C_ receptors, cAMP signals were studied on cells expressed the genetically encoded cAMP sensor Pink Flamindo and recombinant 5-HT_4_ receptors, for monitoring PI3K activity cells stably expressed the genetically encoded PIP_3_ sensor PH(Akt)-Venus were used. It turned out that LY294002 suppressed Ca^2+^ signals initiated by activation 5-HT_2C_ receptors irrespectively of PI3K inhibition, but did not affect cAMP responses initiated by 5-HT_4_ receptors. In turn LY303511 suppressed cAMP signals initiated by 5-HT_4_ receptors, and elicited Ca^2+^ transients exclusively in cells expressed 5-HT_2C_ receptors. Based on these facts and the results of the inhibitory analysis, we hypothesize that the described effects may be due to the activity of LY294002 and LY303511 on the serotonin 5-HT_2C_ and 5-HT_4_ receptors.

## 1. Introduction

The phosphoinositide 3-kinase (PI3K) is lipid kinase phosphorylates phosphatidylinositol (3,4)-bisphosphate (PIP_2_) in the plasma membrane, producing phosphatidylinositol (3,4,5)-trisphosphate (PIP_3_) and stimulating PIP_3_-dependent intracellular processes [1]. PI3K are involved in the regulation of various physiological processes such as cell cycle progression, cell growth, survival and migration, and intracellular vesicular transport [2]. Special attention is paid to the role of PI3K in cancer [3], brain disorders [4], thrombosis and cardiovascular disease [5], and other disease [6].

The role of PI3K in physiological processes can be established by various methods and approaches, but one of the main instruments for such research is PI3K inhibitors [7]. A significant discovery in 1993 was that wortmannin, naturally occurring metabolite of *Penicillium funiculosum*, with anti-inflammatory activities [8], is a highly potent, selective and cell-permeable PI3K inhibitor [9]. Around the same time the first synthetic inhibitor of PI3K LY294002 was generated [10]. Availability of these pharmacological tools, which inhibit all PI3K isoforms, led to an explosion of PI3K research and the identification of many biological processes in which PI3K has a role. On the other hand, it was later shown that these PI3K inhibitors not only to impair activity of PI3Ks from different classes but also to affect unrelated proteins [11]. Notable among them are non-lipid kinases, such as casein kinase-2 (CK2) [12], mammalian target of rapamycin [13], DNA-dependent protein kinase [14], Pim-1 kinase [15], nuclear factor-κB [16, 17], MRP1 [18].

In rat airway smooth muscle cells, wortmannin and LY294002 had opposite effects on serotonin-induced Ca^2+^ signals: while wortmannin caused a small increase in Ca^2+^ responses, LY294002 significantly decreased their magnitudes [19]. Based predominantly on biochemical evidence, the authors inferred that LY294002 affected the serotonin responses independently of PI3K suppression, by acting on, or upstream of, PLC. In our previous study, we also encountered the fact that LY294002 and its structural analogue LY303511 impaired Ca^2+^ signaling induced by serotonin in CHO/5-HT_2C_ cells, the obtained evidence raised the possibility that these compounds could directly affect activity of aminergic GPCRs [20]. Here we studied effects of LY294002 and LY303511 on serotonin induced Ca^2+^ and cAMP signaling in HEK293 cells expressed 5-HT_2C_ or 5-HT_4_ serotonin receptors.

## 2. Materials and methods

### 2.1. Cell lines and culturing

In this study, were employed four monoclonal HEK293 cell lines including HEK/5-HT_2C_, HEK/PF/5-HT_4_, HEK/PF, HEK/PH(Akt)-Venus, and also parental HEK293. Cells were cultured in the Dulbecco’s modified Eagle’s medium (DMEM; Invitrogen) contained 10% (vol/vol) fetal bovine serum (Cytiva), 1% glutamine, 0.1 mg/ml gentamicin (Invitrogen) and selective antibiotic G-418 (0.3 mg/ml) for all monoclonal cells line and additionally Hygromycin B Gold (0.1 mg/ml) for HEK/PF/5-HT_4_ (both from Invivogen). Cells of all mentioned lines were grown in 12-well culture plates at 37 °C in a humidified atmosphere of 5% CO_2_.

### 2.2. Molecular cloning

5-HT_2C_ (GeneBank accession number NM_008312.4) and 5-HT_4_ (GeneBank accession number NM_008313.4) were cloned from mouse brain and subcloned into mammalian expression vectors pDsRed-Monomer-N1 and pAcGFP1-Hyg-N1 respectively as described previously [20, 21]. The presence of DsRed-Monomer or GFP1 allowed for controlling cellular localization of the recombinant protein and easy identification of transfected cells. Pink Flamindo was a gift from Tetsuya Kitaguchi (Addgene plasmid # 102356; http://n2t.net/addgene:102356; RRID:Addgene_102356) [22] and PH(Akt)-Venus was a gift from Narasimhan Gautam (Addgene plasmid #85223; http://n2t.net/addgene:85223; RRID:Addgene_85223) [23].

### 2.3. Monoclonal cell lines

All monoclonal cell lines were derived from HEK293 cells using the same protocol as previously [20, 21]. Before the day of transfection, 3-5 x 10^5^ cells were put in the well of 12-well culture plates in 1 ml of complete growth DMEM medium per well. For transfection of cultured cells, the growth medium was replaced with the transfection mixture, containing 800 μl the growth medium, 200 μl OptiMEM media (Gibco), 2 μl P3000 Reagent (Invitrogen), 2 μl Lipofectamine 3000 (Invitrogen) and 1 μg appropriate expression vector. After 24-hr incubation, the transfection mixture was replaced with the complete growth medium, and transfected cells were cultured in the presence of selective antibiotics for 3 weeks. The antibiotic-resistant cells were analyzed using the FACS Aria SORP sorter (Beckton Dickinson), and those exhibiting most intensive fluorescence at the correct excitation were collected individually for further culturing in a 96-well plate. Next, each monoclone was screened for functionality with Ca2+-, cAMP- and PH(Akt)-Venus-imaging, and monoclones were selected as generating sufficiently high cellular responses to adequate agonists.

As a result, the HEK/PF line stably expressed genetically encoded cAMP sensor Pink Flamindo; HEK/PF/5-HT_4_ stably expressed genetically encoded cAMP sensor Pink Flamindo and recombinant serotonin 5-HT_4_ receptors [21], the HEK/5-HT_2C_ line stably expressed recombinant serotonin 5-HT_2C_ receptors, the HEK/PH(Akt)-Venus line stably expressed PIP_3_ sensor PH(Akt)-Venus.

### 2.4. Ca^2+^ and cAMP imaging

For cell isolation, cultured cells were washed with the Versene solution (Sigma-Aldrich), incubated in the 0.25% Trypsin-EDTA solution (Sigma-Aldrich), resuspended in a complete growth medium. Isolated cells were plated onto a hand-made photometric chamber of nearly 150 μl volume. The chamber was a disposable coverslip (Menzel-Glaser) with an attached ellipsoidal resin wall; the chamber bottom was coated with Cell-Tak (Corning) ensuring sufficient cell adhesion. For Ca^2+^-imaging attached cells were loaded with the Ca^2+^ dye Fura-2 at room temperature (23 – 25 °C) by adding 4 μM Fura-2 AM (AAT Bioquest) and 0.02% Pluronic F-127 (SiChem) to the bath. Cells were loaded for 20 min, rinsed several times, and kept in the bath solution for 1 h prior to recordings. The bath solution contained (mM): 130 NaCl, 5 KCl, 2 CaCl_2_, 1 MgCl_2_, 10 glucose, 10 HEPES (pH 7.4). When necessary, 2 mM CaCl_2_ in the bath was replaced with 0.5 mM EGTA + 0.4 mM CaCl_2_, thus reducing free Ca^2+^ to nearly 260 nM at 23 °c as calculated with the Maxchelator program (http://maxchelator.stanford.edu). The used salts and buffers were from Sigma-Aldrich.

Experiments were carried out using an inverted fluorescent microscope Axiovert 135 equipped with an objective Plan NeoFluar 20x/0.75 (Carl Zeiss), a digital EMCCD camera Luca^EM^ R 604 (Andor Technology), and a hand-made computer-controllable epi-illuminator with a set of light-emitting diodes that enabled multiwave excitation. Fluorescence of the Ca^2+^ dye Fura-2 was excited at 340 ± 6 nm and 380 ± 6 nm, its emission was collected at 520 ± 20 nm. Deviations of cytosolic Ca^2+^ in individual cells are presented as the ratio *F*_340_/*F*_380_, where *F*_340_ and *F*_380_ are the instant intensity of cell fluorescence upon excitation 340 nm and 380 nm, respectively. Fluorescence of cAMP sensor Pink Flamindo was excited at 572 ± 17 nm, its emission was collected at 600 ± 34 nm. Deviations of cytosolic cAMP from the resting level in individual cells were quantified by the ratio Δ*F*/*F*_0_, where Δ*F* = *F* − *F*_0_, *F* is the instant intensity of cell fluorescence, *F*_0_ is the intensity of cell fluorescence recorded in the very beginning of a recording and averaged over a 20 s interval. Serial fluorescent images were captured every second and analyzed using imaging software NIS-Elements (Nikon). All chemicals were applied by the complete replacement of the bath solution in a 150 μl photometric chamber for nearly 2 s using a perfusion system driven by gravity. For data and graphical analysis Sigma Plot 12.5 (Systat Software Inc) was used. “n” represents the number of cells from at least three experiments conducted on different days.

### 2.5. PIP_3_ and Ca^2+^ imaging

HEK/PH(Akt)-Venus cells were cultured in a hand-made photometric chamber for 24 h prior to experiments. The chamber was a Plexiglas framework with an ellipsoidal slot of nearly 150 μl volume, the bottom of the chamber was a disposable coverslip (Menzel-Glaser). Experiments were carried out using an inverted fluorescent microscope Axiovert 200 equipped with an objective Plan NeoFluar 20x/0.75 (Carl Zeiss), a digital sCMOS camera Zyla 4.2P (Andor Technology), metal halide light source AMH-200-F6S (Andor Technology), and spinning disk for confocal microscopy Revolution DSD2 (Andor Technology). Fluorescence of PH(Akt)-Venus was excited at 500 ± 10 nm, emission was collected at 535 ± 15 nm. Serial fluorescent images were captured with NIS-Elements, depending on an experimental protocol. All chemicals were applied by the complete replacement of the bath solution in a 150 μl photometric chamber for nearly 2 s using a perfusion system driven by gravity.

### 2.6. Drugs

The used acetylcholine chloride, insulin, serotonin hydrochloride, RS-102221, and U73122 were purchased from Tocris Bioscience; adenosine and wortmannin were from Sigma-Aldrich; SQ22536 was from Calbiochem; LY294002 hydrochloride, LY303511, were from Tocris Bioscience and MedChemExpress.

## 3. Results

### 3.1. Effects of LY294002 and LY303511 on Ca^2+^ and cAMP signaling initiated by serotonin

Serotonin-induced Ca^2+^ signaling was studied in cells of monoclonal line HEK/5-HT_2C_, stably expressed mouse serotonin 5-HT_2C_ receptor, loaded with the Ca^2+^ dye Fura-2. Monitoring of serotonin-induced cAMP signaling was carried out on cells of monoclonal line HEK/PF/5-HT_4_, stably expressed mouse serotonin 5-HT_4_ receptor and genetically encoded cAMP sensor Pink Flamindo. In a typical experiment, cells were shortly 60 – 100 s stimulated by serotonin at moderate concentrations every 300 – 900 s, the period being sufficient for post-stimulation recovery of cells and prolongation of their responsiveness during long-lasting recordings. The pulse stimulation of HEK/5-HT_2C_ cells with serotonin triggered Ca^2+^ transients in control but not in the presence of PI3K inhibitor LY294002, while applications of LY303511, inactive analogue of LY294002, triggered agonist-like Ca^2+^ transients (Fig. 1A). In turn, magnitude of cAMP signals elicited by serotonin in HEK/PF/5-HT_4_ cells decreased in the presence of LY303511, whereas LY294002 did not affect these responses (Fig. 1B).

**Fig. 1.**
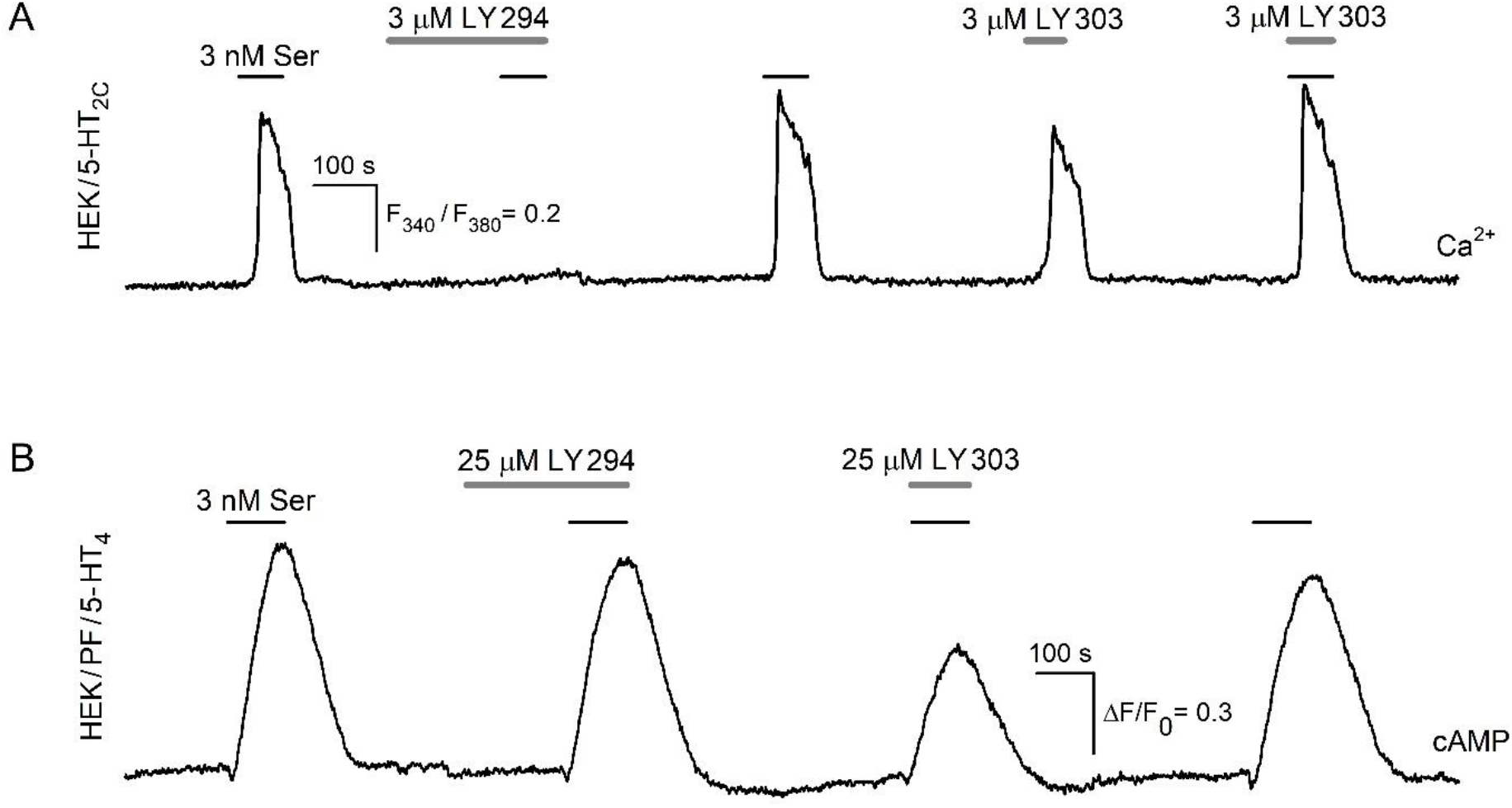
Effects of LY294002 and LY303511 on Ca^2+^ and cAMP signaling. Representative recordings of intracellular (A) Ca^2+^ in individual Fura-2-loaded HEK/5-HT_2C_ cell and (B) cAMP in individual HEK/PF/5-HT_4_ cell stimulated by serotonin in control and in the presence of LY294002 or LY303511. (A) LY294002 (3 μM) blocked Ca^2+^ signals elicited by serotonin (3 nM), while LY303511 (3 μM) induced Ca^2+^ transients and did not inhibit cell responsiveness to serotonin. (B) LY294002 (25 μM) did not inhibit cAMP signals induced by 3 nM serotonin, while LY303511 (25 μM) decreased their magnitude. Here and in the below figures: applications of compounds are indicated by the straight-line segments above the experimental traces. Deviations of cytosolic Ca^2+^ in individual Fura-2-loaded cells are presented as the ratio *F*_340_/*F*_380_, where *F*_340_ and *F*_380_ are the instant intensity of cell fluorescence upon excitation 340 nm and 380 nm, respectively. Deviations of cytosolic cAMP in individual cells expressed Pink Flamindo were quantified by the ratio Δ*F*/*F*_0_, where Δ*F* = *F* − *F*_0_, *F* is the instant intensity of cell fluorescence, *F*_0_ is the intensity of cell fluorescence recorded in the very beginning of a recording and averaged over a 20 s interval.

### 3.2. Effects of LY294002 and LY303511 on PI3K activity

The described effects of PI3K inhibitor LY294002 and its inactive analogue LY303511 on cell responsiveness to serotonin (Fig. 1) were difficult to explain in terms of their claimed activity on PI3K. We therefore attempted to evaluate specific activity of LY294002 and LY303511 on PI3K by using HEK/PH(Akt)-Venus cells, which stably expressed genetically encoded PIP_3_ sensor PH(Akt)-Venus. The total fluorescence of a PH(Akt)-Venus expressing cell is apparently independent of a PIP_3_ level, while generation of PIP_3_ in the plasmalemma stimulates translocation of PH(Akt)-Venus from the cytosol to plasma membrane [23]. To initiate PIP_3_ generation, we used insulin known to stimulate tyrosine kinase receptors coupled particularly to the PI3K/Akt pathway [24]. Based on this and using the protocol shown on Fig. 2A, we examined the effects of LY294002 and LY303511 on PI3K. It turned out that 30 μM LY294002 prevented the insulin-induced translocation of PH(Akt)-Venus to the plasmalemma (Fig. 2B, middle panel). When cells were rinsed out for nearly 10 min to remove this PI3K inhibitor, insulin applied once again was capable of initiating the PH(Akt)-Venus translocation, which was indicated by the appearance of relatively bright and contrast narrow zones corresponding to plasma membranes (Fig. 2B, right panel). In the similar assay, LY303511 at the same dose did not prevent insulin induced translocation of PH(Akt)-Venus to the plasmalemma (Fig. 2C, middle panel). These observations indicated that LY294002 indeed effectively inhibited PI3K, whereas LY303 had no effect on its function.

**Fig. 2.**
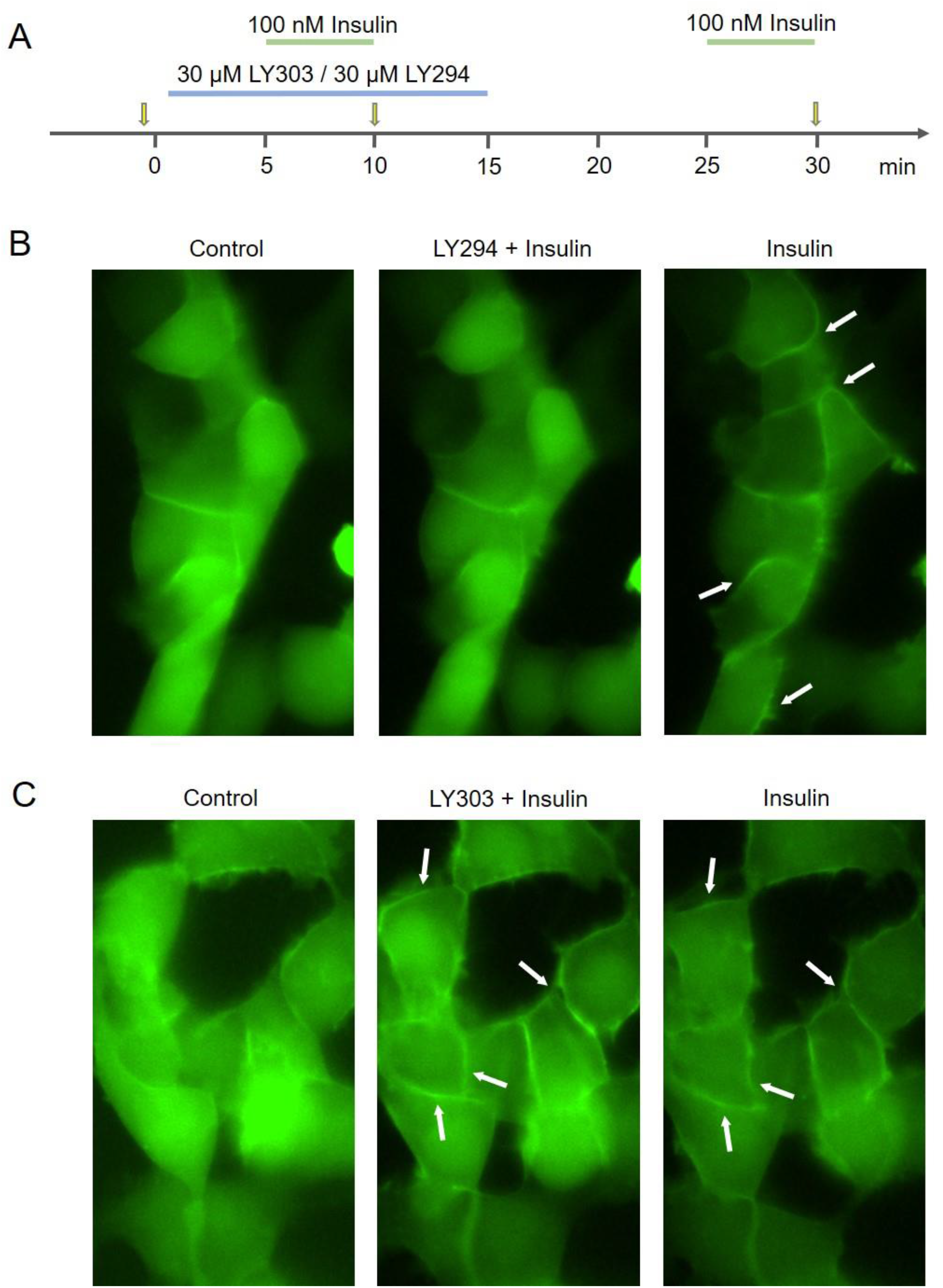
Activity of LY294002 and LY303511 against PI3K. (A) Experimental protocol timing. The applications of LY294002 or LY303511, and insulin during recordings are indicated by the line segments above the time axis; the moments of image capturing are indicated by the yellow arrows. (B, C) Representative sequential images of HEK/PH(Akt)-Venus cells obtained at the time points indicated in (A). The left panels, control images of the group of cells obtained right before drug application demonstrate the virtually homogeneous distribution of PH(Akt)-Venus fluorescence over cell bodies. The middle panels, the cell stimulation with insulin (100 nM) in the presence of LY294002 (30 μM) negligibly affected sensor distribution (B), while in the presence of LY303511 (30 μM) insulin initiated a drop in the fluorescence of the cell cytosol and the appearance of detectable stripe-like fluorescent zones indicated by the white arrows (C), which suggesting the insulin-induced accumulation of the PH(Akt)-Venus sensor in the plasmalemma. The right panels, images obtained after the rinse of LY294002 or LY303511 and stimulation with insulin. As shown, LY303511 negligibly affected insulin-induced PI3K activity, while LY294002 inhibited it.

### 3.2. Cellular target of LY303511

Thus, we are convinced that the effect of inhibition of serotonin-induced cAMP signals (Fig. 1B) in the presence of LY303511 is not associated with inhibition of PI3K (Fig. 2C), and indicates that LY303511 acts on a cellular target other than PI3K. Particularly intriguing was the ability of LY303511 induce Ca^2+^ transients in HEK/5-HT_2C_ cells. We attempted to characterize these LY303511-induced Ca^2+^ responses and found that the reduction of bath Ca^2+^ from 2 mM to 260 nM weakly or negligibly affected these signals, which completely disappeared in the presence of PLC inhibitor U73122 (Fig. 3A). These observations indicated that LY303511-induced Ca^2+^ transients are generated through the phosphoinositide pathway and suggested that these signals could be triggered by the GPCRs. In this regard, it is critical to note that LY303511 did not stimulate Ca^2+^ transients in parental HEK293 cells, which not expressing recombinant 5-HT_2C_ receptors (Fig. 3B). This suggests the possibility that LY303511 specifically activated 5-HT_2C_ receptors, leading to the generation of Ca^2+^ transients through the phosphoinositide pathway. This idea was supported by the fact that Ca^2+^ responses of HEK/5-HT_2C_ cells to LY303511 were invariably blocked in the presence of the specific 5-HT_2C_ receptor antagonist RS-102221 (Fig. 3C).

**Fig. 3.**
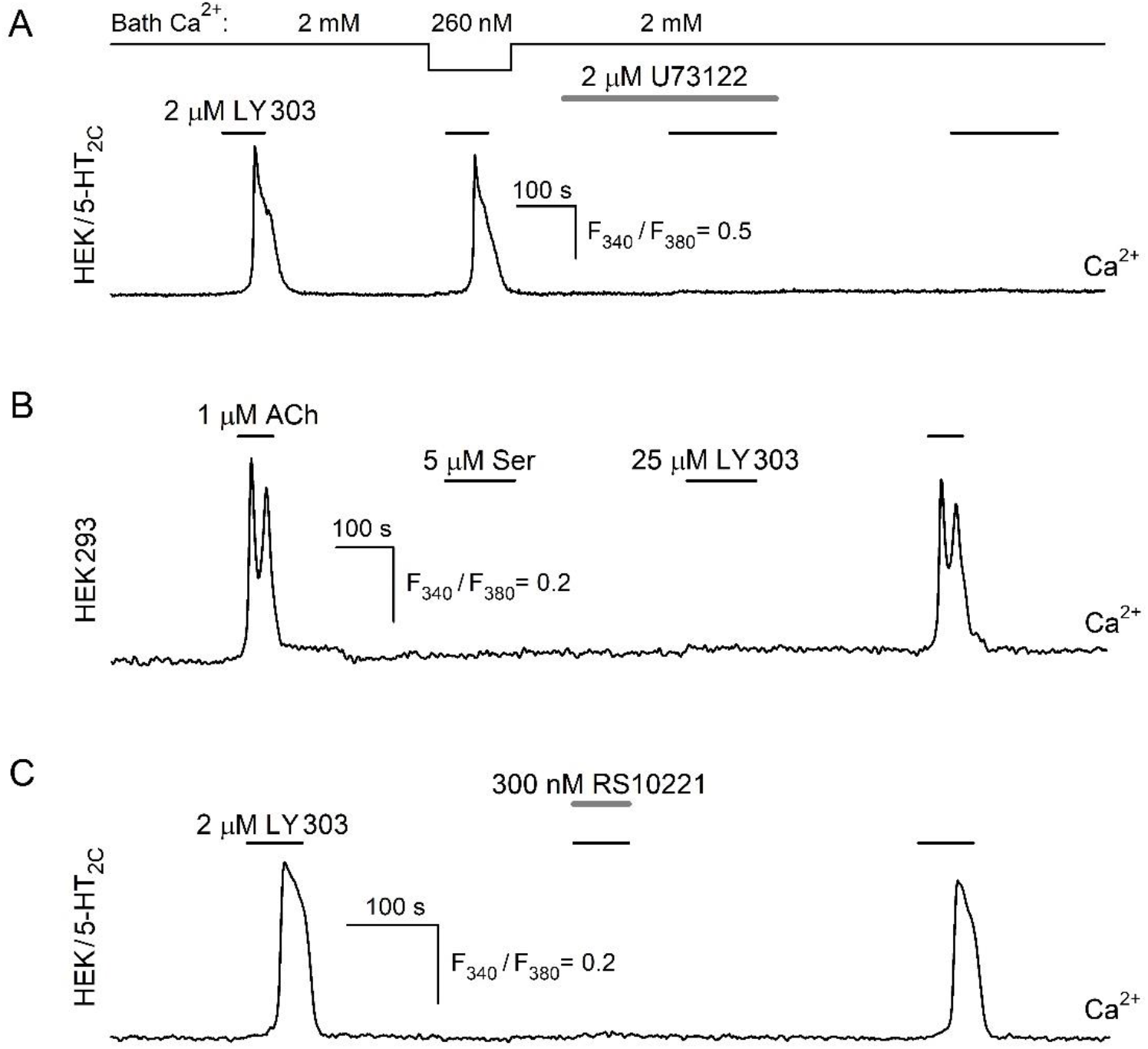
Evidence for LY303511 activates serotonin 5-HT_2C_ receptors. (A) Ca^2+^ signals elicited by LY303511 (2 μM) in HEK/5-HT_2C_ cells were weakly affected by the reduction of bath Ca^2+^ from 2 mM to 260 nM and suppressed by the PLC inhibitor U73122 (2 μM). (B) Parental HEK293 cells were not sensitive to serotonin as well as LY303511, the ability of cells to generate Ca^2+^ transients were tested by stimulation with acetylcholine (ACh) (1 μM). (C) Ca^2+^ signals initiated in HEK/5-HT_2C_ cells by LY303511 (2 μM) were blocked in the presence of 5-HT_2C_ receptor antagonist RS102221 (300 nM).

### 3.3. Cellular target of LY294002 unrelated to PI3K

The inhibitory effects of LY294002 on Ca^2+^ responses HEK/5-HT_2C_ cells to serotonin (Fig. 1A) could be considered as evidence for the involvement of PI3K in the transduction of this agonist, but another PI3K inhibitor wortmannin (1 – 10 μM) did not affect the responsiveness of the same cells to serotonin (Fig. 4A). LY294002 is also known to block CK2 [12], and we tested whether this was responsible for the inhibitory effect of this compound. The CK2 inhibitor hematein did not affect the ability of cells to generate Ca^2+^ responses to serotonin, while LY294002 completely blocked them (Fig. 4B). These findings shown that observed inhibitory effect of LY294002 on serotonin-induced Ca^2+^ transients in HEK/5-HT_2C_ cells was not specifically associated with the suppression of PI3K or CK2 activity.

**Fig. 4.**
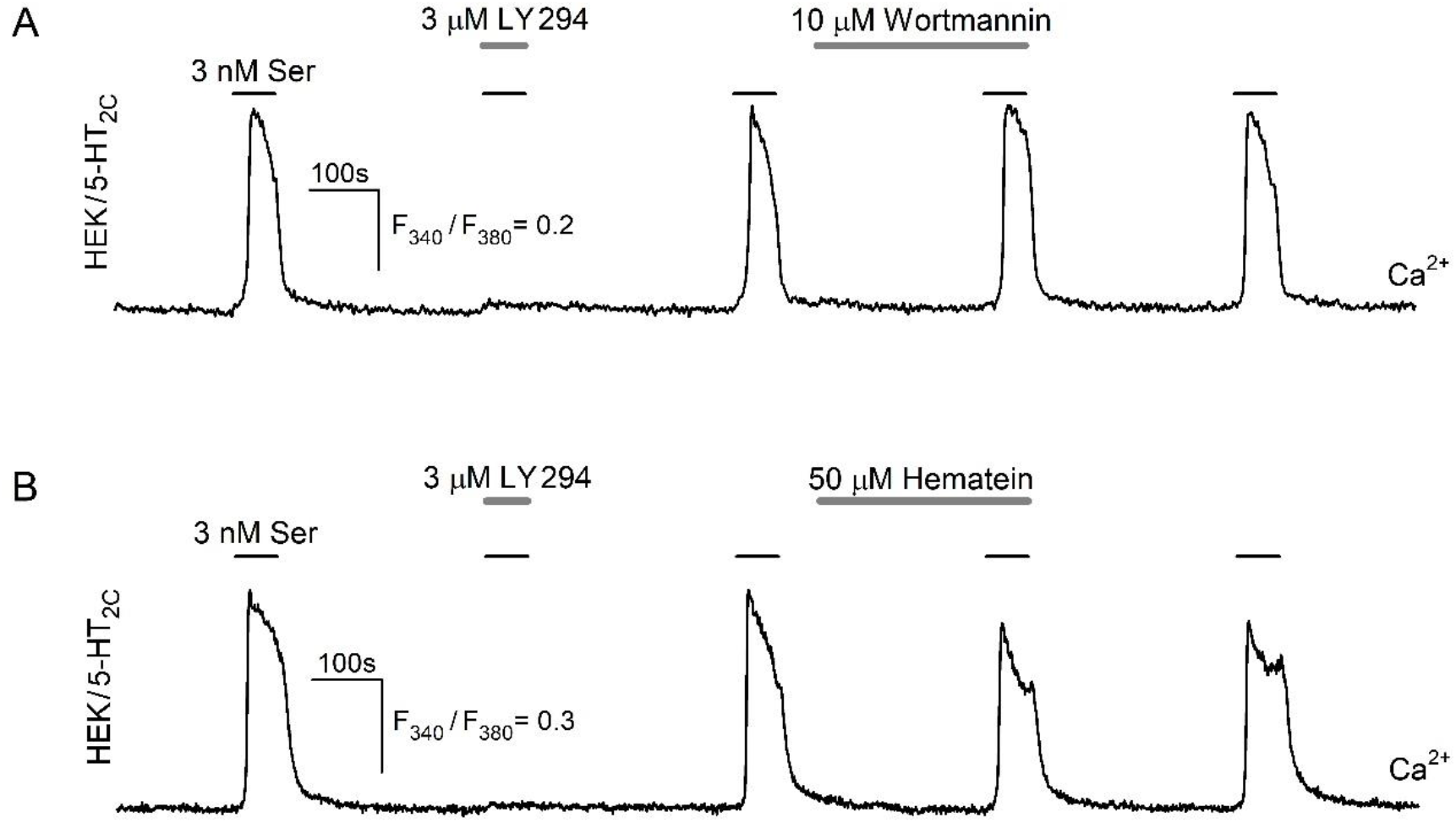
LY294002 blocks serotonin-induced Ca^2+^ transients irrespectively of PI3K or CK2 inhibition. Representative recordings demonstrating dissimilar effects of LY294002 (3 μM) and inhibitor PI3K wortmannin (10 μM) (A) or CK2 inhibitor hematein (50 μM) (B) on serotonin-induced Ca^2+^ transients in HEK/5-HT_2C_ cells.

In an attempt to identify the cellular target of LY294002 and LY303511, we compared the inhibitory effects of these compounds on Ca^2+^ or cAMP signals induced by serotonin and another agonist elicited similar signals. Serotonin-induced Ca^2+^ transients were completely blocked in the presence of the PLC inhibitor U73122 (Fig. 5A), in its turn cAMP signals were inhibited by the AC inhibitor SQ22536 (Fig. 5B). Ca^2+^ and cAMP responses initiated by P1 receptors agonist adenosine had the same inhibitory profile (Fig. 5C,D), so we tested how LY294002 or LY303511 affected them. It turned out that LY294002 and LY303511 negligibly affected transduction of adenosine (Fig. 5E). It means the PLC or AC pathway could hardly be the target of these compounds, otherwise, LY294002 and LY303511 would detectably influence Ca^2+^ or cAMP responses to adenosine, as was the case with serotonin (Fig. 1). The peculiar feature of the LY294002 and LY303511 effects was that these compounds prevented Ca^2+^ or cAMP signals even if they were applied simultaneously with the serotonin (Fig. 1). The inhibition of intracellular enzymes by an extracellularly applied substance should have occurred with a certain delay because it would take some time for penetrating through the plasma membrane and being accumulated in the cytosol at an appropriate level. The combination of the fact that LY294002 and LY303511 did not affect signaling molecules downstream of the PLC and AC pathway, with the fact that LY294002 and LY303511 were capable of suppressing Ca^2+^ or cAMP responses to serotonin without preincubation pointed out the possibility that these chemicals acted extracellularly.

**Fig. 5.**
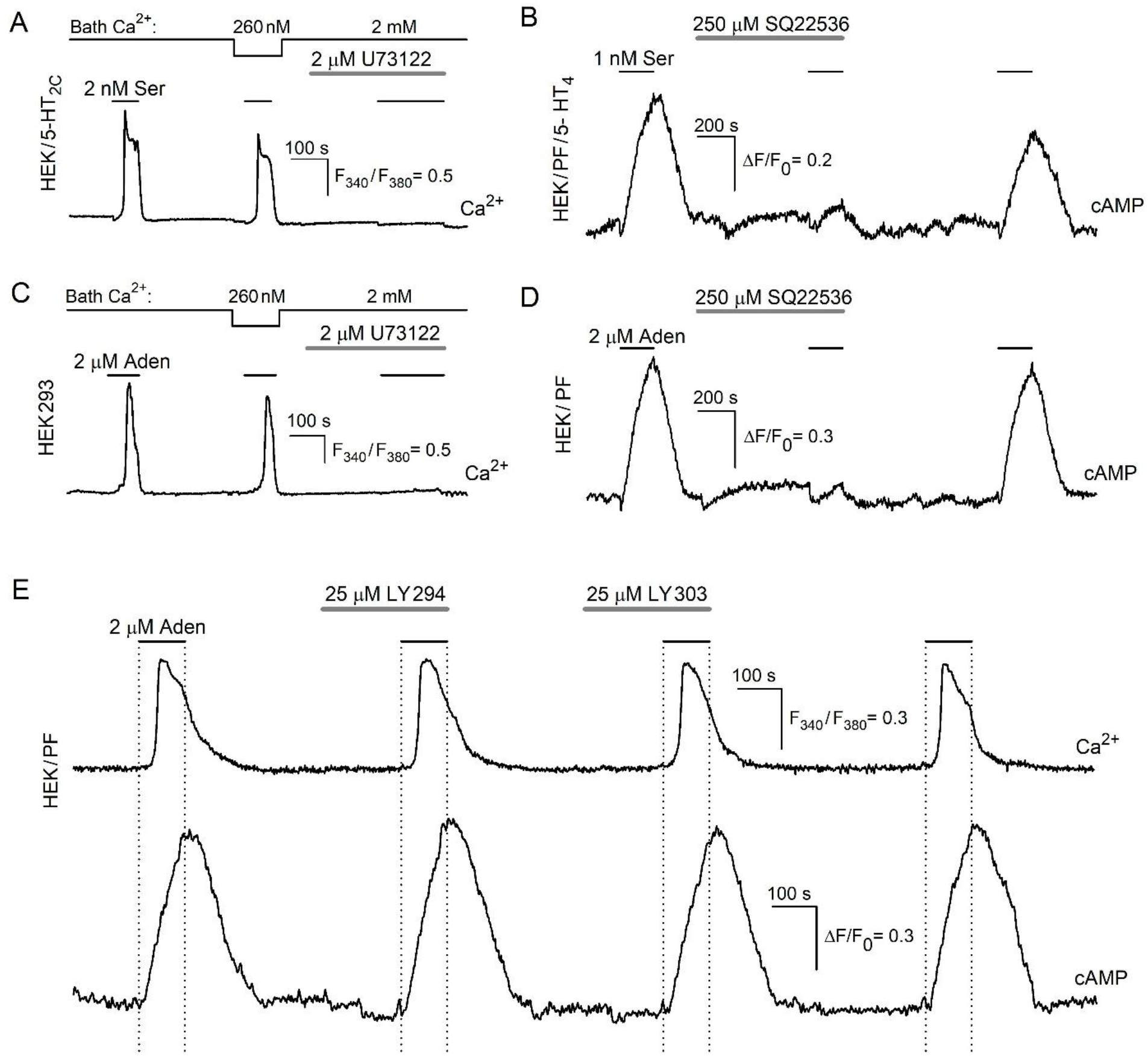
Comparison of Ca^2+^ and cAMP responses to serotonin and adenosine. (A – D) Representative recordings demonstrating the involvement of the phosphoinositide cascade in Ca^2+^ transients and AC cascade in cAMP signals elicited by serotonin or adenosine. Serotonin-induced Ca^2+^ transients in HEK/5-HT_2C_ cells were weakly affected by the reduction of bath Ca^2+^ from 2 mM to 260 nM and suppressed by the PLC inhibitor U73122 (2 μM) (A), while cAMP signals in HEK/PF/5-HT_4_ cells were inhibited in the presence of AC inhibitor SQ22536 (250 μM) (B). Adenosine-initiated Ca^2+^ and cAMP signals were tested on HEK293 and HEK/PF cells, respectively, and demonstrated the similar inhibitory profile (C, D) to the signals initiated by serotonin (A, B). (E) Representative recording of simultaneous intracellular Ca^2+^ and cAMP monitoring in a single HEK/PF cell demonstrating no effect of LY294002 (25 μM) or LY303511 (25 μM) on Ca^2+^ and cAMP signaling initiated by adenosine (2 μM).

## 4. Discussion

The first identified PI3K inhibitors were wortmannin, a naturally occurring metabolite of *Penicillium funiculosum* [9, 25] and the synthetic compound LY294002, which was generated together with its inactive structural analogue LY303511 [10] (Fig. 6). Being considered as rather specific PI3K inhibitors for a long time, wortmannin and LY294002 were widely used to verify a role for the PI3K pathway in cellular functions, while LY303511 as a control was and is still rarely used. However, these PI3K inhibitors were later shown not only to impair activity of PI3Ks from different classes but also to affect unrelated proteins involved in various physiological processes, particularly, metabolism, transcription or protein trafficking and dynamics [11]. Notable among them are non-lipid kinases, such as mammalian target of rapamycin [13], DNA-dependent protein kinase [14], casein kinase-2 (CK2) [12], Pim-1 kinase [15], nuclear factor-κB [16, 17], MRP1 [26].

**Fig. 6.**
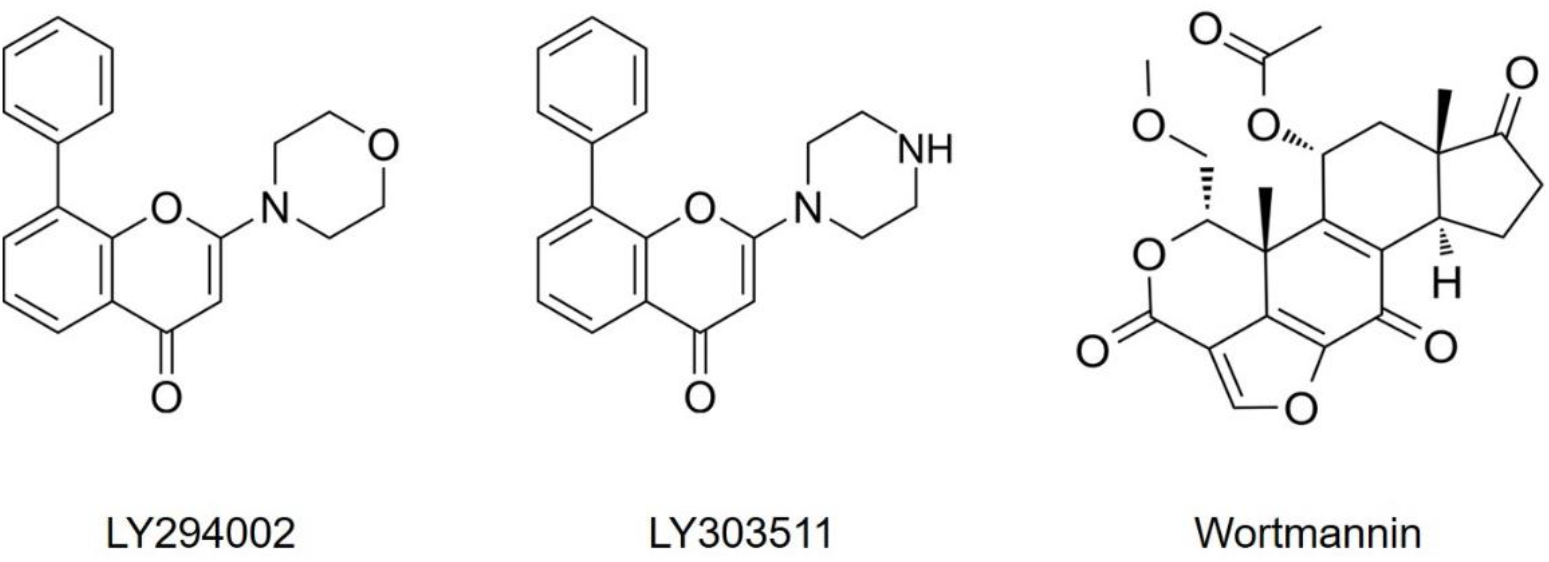
Structural formulas of LY294002, LY303511, and wortmannin.

LY294002 has been demonstrated to affect serotonin-induced Ca^2+^ signaling in airway smooth muscle cells independently of PI3K inhibition [19]. Based predominantly on biochemical evidence, the authors inferred that LY294002 affected the serotonin responses independently of PI3K suppression, by acting on, or upstream of, PLC. In our previous study, we also encountered the fact that LY294002 impaired Ca^2+^ signaling induced by serotonin in CHO/5-HT_2C_ cells, the obtained evidence raised the possibility that these compounds could directly affect activity of GPCRs [20]. Here we focused on studying effects of LY294002 and LY303511 on Ca^2+^ signaling in HEK293 cells expressed serotonin 5-HT_2C_ receptors and for the first time showed the effects of these compounds on serotonin-induced cAMP signaling on cells expressed serotonin 5-HT_4_ receptors. We showed that LY294002 blocks serotonin-induced Ca^2+^ transients, independent of PI3K and CK2 inhibition (Fig. 4). In turn, LY303511, inactive against PI3K, decreased the magnitudes of cAMP signals elicited by serotonin (Fig. 1). The above clearly indicates that LY303511, like LY294002, has cellular targets other than PI3K. The most striking effect of LY303511 was that this compound triggered agonist-like Ca^2+^ transients in cells expressing serotonin 5-HT_2C_ receptors, while no similar effect was observed in cells without this receptor (Fig. 3). This fact and the results of the inhibitory analysis directly indicate that the target of LY303511 was precisely the serotonin 5-HT_2C_ receptors, which it activated (Fig. 3). The comparison results of the effects LY294002 and LY303511 on signals initiated by serotonin and adenosine (Figs. 1 and 5E) indicated the possibility of these compounds acting directly on 5-HT_2C_ and 5-HT_4_ serotonin receptors, respectively.

There are many studies where LY294002 was used to investigate various processes triggered by serotonin [27-30]. Only a few of them used another PI3K inhibitor, wortmannin, to confirm the obtained results [31, 32]. It is important to note that we were unable to find a single study where the effects of LY294002 on serotonin-induced processes were tested using the inactive analogue of this compound, LY303511. An example of such work is a study where Tourette syndrome was induced by the selective 5-HT_2A/2C_ agonist, and LY294002 significantly reduced all detected effects of this model. The authors concluded that the protective effect of LY294002 against Tourette syndrome might be associated with the regulation of PI3K/Akt/NF-B pathway, although the findings were not confirmed with any other PI3K inhibitor or LY303511. Based on the results of the present study, it could be assumed that in that case the effects of LY294002 could be associated not with inhibition of PI3K, but with its inhibitory effect on 5-HT_2C_ receptors, which led to impaired induction of Tourette Syndrome [33].

In this study we showed that LY294002 suppressed Ca^2+^ signaling initiated by activation serotonin 5-HT_2C_ receptors irrespectively of PI3K inhibition, but did not affect cAMP responses initiated by serotonin 5-HT_4_ receptors. In turn PI3K-inactive structural analogue of LY294002, LY303511 suppressed cAMP signals initiated by serotonin 5-HT_4_ receptors, and induced Ca^2+^ transients in 5-HT_2C_ expressed cells only. Based on the results of the inhibitory assay, we hypothesize that the described effects may be due to the acting of LY294002 and LY303511 directly on the 5-HT_2C_ and 5-HT_4_ serotonin receptors. Thus, to demonstrate the contribution of PI3K to serotonin-induced processes, LY294002 and LY303511 should be used with caution, and results obtained with these compounds should be confirmed using PI3K inhibitors with significantly different chemical structures, such as wortmannin.

## Funding

This work was supported by the Russian Science Foundation [grant 19-75-10068].

## CRediT authorship contribution statement

**Polina D. Kotova:** Supervision, Investigation, Visualization, Writing – original draft, Funding acquisition. **Ekaterina A. Dymova:** Investigation, Visualization. **Olga A. Rogachevskaja:** Investigation. **Stanislav S. Kolesnikov:** Conceptualization, Writing – review & editing

## Declaration of competing interest

The authors declare that they have no known competing financial interests or personal relationships that could have appeared to influence the work reported in this paper.

## Acknowledgements

The authors thank Daria M. Potashnikova for her assistance in sorting cells on the FACSAria SORP cell sorter (MSU Program of Development).

## Notes

### Competing Interest Statement

The authors have declared no competing interest.

